# Long-read metagenomics reveals unexplored viral diversity and genomic novelty in the mesopelagic Andaman Sea

**DOI:** 10.64898/2026.11.18.764573

**Authors:** Manisha Ray, Muthuvarmadam Subramanian Ram, Himanshu Kothari, Mamidala Srilakshmi, Krishnan Harinivas Harshan, Govindhaswamy Umapathy

## Abstract

The mesopelagic zone (200–1000 m) of the ocean represents Earth’s largest yet most underexplored biome, particularly with respect to viral diversity and function. The deep-sea virome of the Indian Ocean remains largely unexplored despite the ecological significance of marine viruses in regulating microbial mortality, biogeochemical cycling and carbon flux via the viral shunt. In this study, we explored the diversity of DNA viruses across mesopelagic depths in the Andaman Sea. Seawater samples collected from multiple depths were enriched for viral particles, followed by viral DNA extraction and long-read metagenomic sequencing using Oxford Nanopore Technologies. We used three complementary approaches for the identification of assembled contigs of viruses. Based on the consensus from these three approaches followed by quality assessment, we retained 1,604 viral contigs of high-confidence and high-quality. Among the taxonomically assigned viruses, members of the class *Caudoviricetes* dominated across all samples. The assignment decreased substantially at lower taxonomic ranks, highlighting the limited representation of environmental viruses in existing reference databases. Gene-sharing network analysis further revealed genomic novelty and showing limited gene-sharing relationships with reference genomes. Host prediction analyses assigned putative hosts to 10.7% of viral contigs, of which 98.3% were associated with bacteria and 1.7% with archaea. Among predicted bacterial hosts, Pseudomonadota predominated. Structure-informed functional annotation of complete viral contigs identified predominantly structural and tail-associated proteins, while a substantial proportion of viral proteins remained functionally uncharacterized. Collectively, these findings reveal previously under-explored viral diversity in the mesopelagic Andaman Sea study and provide the first comprehensive insight into the deep-ocean virome of this region.

## Introduction

The deep ocean, typically defined as waters below 200 m, represents the largest and least explored biome on Earth, accounting for over 95% of the total ocean volume and approximately 65% of the planet’s surface area [1]. This vast realm is subdivided into the mesopelagic (200–1,000 m), bathypelagic (1,000–4,000 m), and abyssopelagic zones (>4,000 m), each characterized by a progressive absence of solar radiation, extreme hydrostatic pressure, low temperatures and diminishing nutrient flux from the photic zone above [2]. Despite constituting the dominant fraction of the ocean’s habitable space, the deep sea remains among the most poorly sampled environments on Earth [1]. This persistent undersampling reflects logistical and technical constraints [3]. The Intergovernmental Panel on Climate Change (IPCC) has acknowledged the deep ocean as severely underrepresented in global climate and ecosystem assessments, in carbon storage dynamics, and in vulnerability to ongoing anthropogenic stressors, such as global warming, deoxygenation, and acidification [4]. Far from being a biologically inert abyss, the deep ocean is a key engine of global biogeochemical cycling. The organisms that drive these transformations are bacteria, archaea, protists, and the viruses that infect them, which remain incompletely catalogued, limiting our comprehension of global biogeochemical models.

Viruses constitute the most abundant and genetically diverse biological entities in the marine biosphere, with estimated oceanic abundances approaching 10^30^ particles [3, 5]. Through lytic infection, viruses regulate microbial population dynamics and mediate the release of cellular contents into dissolved organic matter pools via the “viral shunt,” thereby impacting carbon and nutrient fluxes [3]. In addition, viruses encode auxiliary metabolic genes (AMGs), host-derived genes retained in viral genomes that are expressed during infection to rewire host metabolism toward viral replication [6]. AMGs encoding genes involved in photosynthesis, carbon fixation, phosphate acquisition, nitrogen cycling and sulfur metabolism have been catalogued across oceanic viromes [6–8].

Over the past decade, large-scale efforts such as the Tara Oceans expedition and the Global Ocean Virome (GOV) project have significantly expanded our understanding of marine viral diversity, revealing vast viral richness and previously unknown viral lineages [9, 10]. However, these studies have predominantly focused on surface and epipelagic waters, with comparatively limited representation of deeper ocean layers. Furthermore, geographic biases persist, with underrepresentation of regions such as the Indian Ocean and, in particular, the Andaman Sea. As a result, the viral diversity and ecological roles of viruses in these underexplored regions remain largely unknown.

The paucity of deep-ocean viromics data is not solely due to limited sampling efforts, but also reflects genuine technical bottlenecks that are magnified at depth. Viral particles in the deep ocean are present at markedly lower concentrations than in surface waters, often falling below 10^5^ particles mL^−1^ at mesopelagic and bathypelagic depths, compared with 10^6^–10^7^ particles mL^−1^ in surface waters [5, 11]. Generating viromes from these low-biomass environments requires efficient concentration of viral particles from large water volumes (typically 20–200 L) using tangential flow filtration (TFF), ultracentrifugation, iron chloride flocculation or combinations thereof, each method introducing its own biases and losses [12–14]. Density gradient ultracentrifugation, while effective for purification, is time-consuming, requires specialized instrumentation, and can result in substantial viral loss [15]. Traditional short-read (150-300 bp) sequencing approaches, while widely used, often yield fragmented assemblies that hinder accurate genome reconstruction and functional annotation [16]. In contrast, long reads can span entire viral genomes in a single read, substantially reducing assembly fragmentation and recovering viral genomic sequences that remain unresolved in short-read datasets [17].

Beyond technical sampling constraints, the bioinformatic analysis of viral metagenomes presents distinct conceptual challenges that distinguish it from bacterial and archaeal metagenomics. Unlike cellular organisms, for which the 16S rRNA gene provides a universal phylogenetic marker enabling culture-independent identification, viruses lack an equivalent conserved marker gene spanning all viral lineages [3, 18, 19]. Viral taxonomy, therefore, relies on whole-genome comparisons, protein cluster analyses and hallmark gene phylogenies. These methods are computationally intensive and inherently dependent on the completeness of reference databases [20, 21]. But a substantial fraction of viral diversity, often referred to as “viral dark matter”, remains uncharacterized due to limited representation in reference databases [18]. This limitation constrains both taxonomic assignment and functional annotation, particularly in understudied environments such as the deep ocean.

While research has begun to characterise the surface microbiome of Andaman coastal waters [22], no study has examined the virome of this region at any depth of the mesopelagic zone. Here, we present the first viromics study of this region, recovering DNA viral metagenomes across a depth gradient spanning the mesopelagic zone. We specifically aimed to: (i) reconstructing the high-quality and high-confidence viral contigs from the mesopelagic zone of the Andaman Sea; (ii) decipher the identity of the DNA viruses at depth in the Indian Ocean and (iii) delineate the novelty of these reconstructed viral contigs, their potential roles and hosts.

## Methods

### Study region, sampling and abiotic data

Seawater samples were collected from different depths (200 m, 500 m and 900 m) in the Andaman Sea using a CTD (conductivity–temperature–depth) rosette system fitted with Niskin sampling bottles on FORV Sagar Sampada 412 cruise. The rosette assembly was deployed to the target depths, and 10 L of seawater was collected from each depth layer. Simultaneously, physicochemical parameters including temperature, salinity, dissolved oxygen, and depth were measured using onboard CTD sensors. The detailed table of all measured abiotic variables, geo-coordinates (Figure 1), and other accessory information is provided in Supplementary 1.

**Figure 1:**
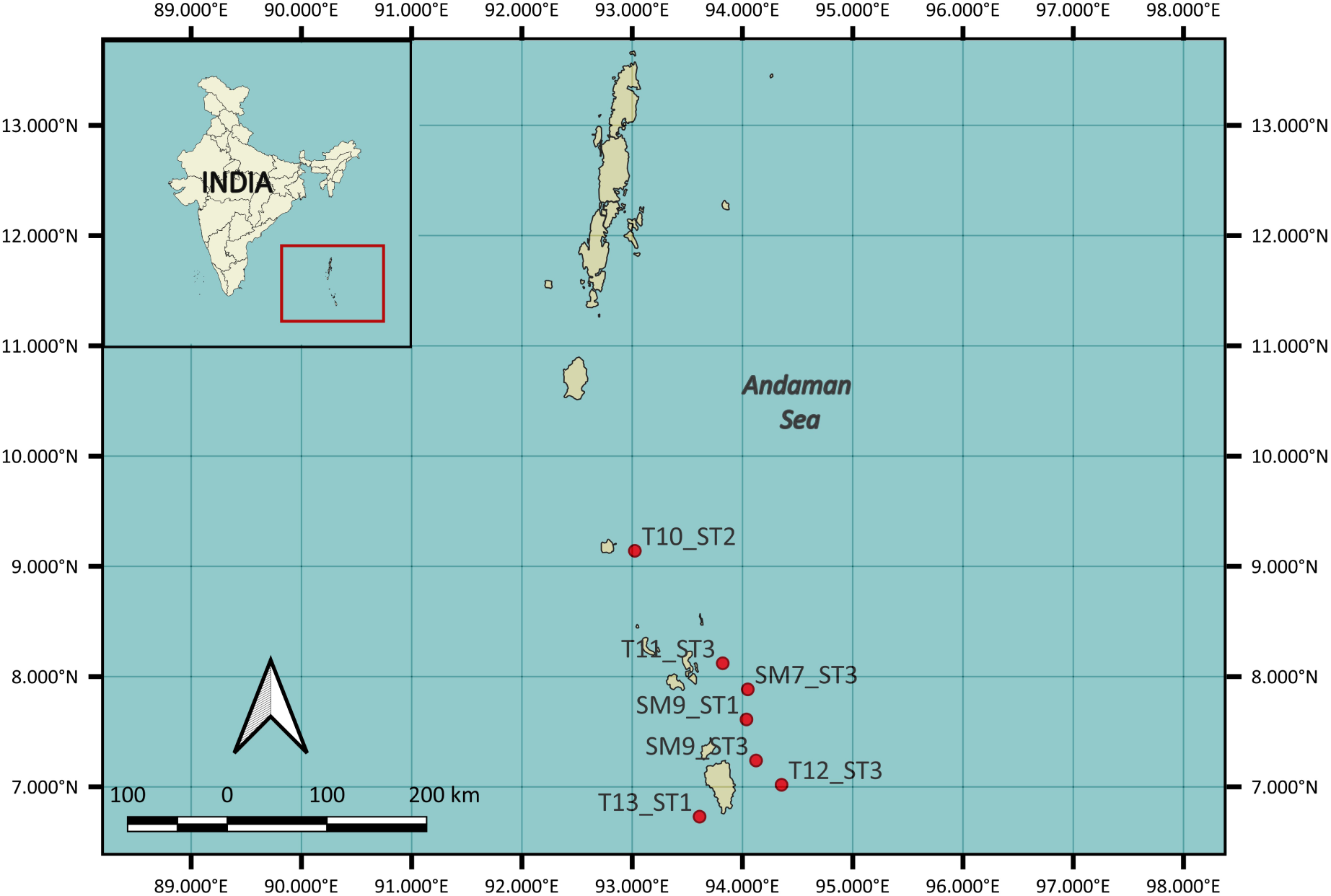
Map depicting the sampled locations from Andaman Sea. Sampling was conducted using a CTD rosette system equipped with Niskin bottles on FORV Sagar Sampada 412 cruise to collect seawater from mesopelagic depths ranging from 200 m to 900 m. The seamounts are labelled as “SM” and transects as “T”. The samples are from various depths of mesopelagic zone ranging from 200 meters to 900 meters.

Hydrographic data were obtained from CTD (Sea-Bird Electronics) profiles and processed in R (version 4.4.1) using the tidyverse v.2.0.0. The calibrated .cnv files were parsed to extract key environmental variables, including temperature, salinity, dissolved oxygen, fluorescence, photosynthetically active radiation (PAR) and conductivity, along with latitude and longitude.

Because each biological sample corresponded to a unique depth, environmental variables were matched to the actual sampling depth (200 m, 500 m, or 900 m) rather than averaged across the water column. For each CTD profile, the measurement closest to the target depth or for the same depth was selected, ensuring an accurate representation of environmental conditions. The final environmental dataset was used for downstream multivariate statistical analysis as mentioned in section 2.9.

### Enrichment of viral particles

The process of enriching viral particles from seawater consisted of three major steps: pre-filtration, concentration, and removal of free nucleic acids from water samples.

Seawater samples were subjected to sequential filtration to remove particulate matter and cellular organisms before viral concentration. Initially, samples were passed through a 3 µm pore-size polycarbonate (PC) filter to eliminate large debris and planktonic material. The filtrate was subsequently passed through a 0.22 µm pore-size PC filter to remove bacteria, thereby enriching the filtrate for viral particles. Viral particles in the filtrate were concentrated using an iron chloride (FeCl₃) based flocculation approach [23]. Briefly, 2 mL of 100 g L⁻¹ Fe stock solution was added to 10 L of filtrate (final concentration: 10 mg L⁻¹ Fe), followed by gentle mixing and incubation at room temperature for 1 h to facilitate floc formation. The Fe-virus flocs were then collected by filtration through 1.2 µm pore-size PC or PES filters. Filters containing Fe-virus flocs were stored in the dark at 4°C until further processing in the lab. Viral particles were subsequently resuspended from the filters using 10 mL of ascorbate-EDTA buffer per filter, with gentle agitation using a sterile L-shaped spreader. The resuspended viral fraction was further concentrated using 100 kDa centrifugal ultrafiltration units (Amicon® Ultra MilliporeSigma, Burlington, MA, USA), reducing the volume to approximately 1.5–2 mL. To ensure that only encapsidated viral nucleic acids were retained, concentrated viral suspensions were treated with DNase and RNase according to the manufacturer’s instructions. This step enabled the degradation of extracellular DNA and RNA. Enzymatic activity was subsequently terminated by EDTA before downstream viral DNA extraction.

### Transmission electron microscopic observation

The Amicon® centrifuged samples were used to visualise the viral particles using transmission electron microscope. The phage virions were allowed to adsorb to the formvar-coated copper grid for 30 minutes. Following this, a negative staining was performed with uranyless at room temperature by incubating for 5 minutes. The grid was then air-dried at room temperature before loading and observing for TEM. The micrographs were taken in Thermo Scientific Talos L120C 120 kV TEM.

### Isolation of viral DNA

Viral DNA was extracted from concentrated viral suspensions using AllPrep PowerViral DNA/RNA Kit, (QIAGEN, Germany) with minor modifications to optimize recovery from purified viral fractions. As viral particles had already been enriched and separated from cellular material, the bead-beating step using glass beads was omitted to avoid DNA shearing. The extraction protocol was scaled up according to the input volume of viral concentrate. Samples were processed through the silica-based spin columns following the manufacturer’s instructions for binding, washing and purification steps. Viral DNA was eluted in 30 µL of elution buffer and stored at −20°C until further use. The purity of extracted DNA was measured using a NanoDrop spectrophotometer, based on absorbance ratios (A260/A280 and A260/A230). DNA concentration was quantified with a Qubit Fluorometer, following the manufacturer’s protocol. Only samples meeting quality and concentration thresholds were used for downstream sequencing.

### Preparation of library and its sequencing

The extracted DNA was used as input for the rapid PCR Barcoding Kit SQK-RPB114.24 (Oxford Nanopore Technologies, Oxford, UK) protocol for library preparation. Briefly, approximately 3 ng of input DNA was used for library preparation. DNA samples were amplified and barcoded using the rapid PCR barcoding kit to enable multiplexed sequencing following the manufacturer’s protocol. Briefly, DNA was subjected to transposase-mediated fragmentation and adapter ligation, followed by PCR amplification to incorporate barcodes. Libraries were pooled in equimolar concentrations and loaded onto an Oxford Nanopore R10.4.1 flow cell. Sequencing was performed on the GridION platform using standard MinKNOW settings. Basecalling and demultiplexing were performed using ONT’s Dorado v.0.8.2 basecaller and model dna_r10.4.1_e8.2_400bps_sup@5.0.0 with default settings.

### Quality filtering and assembly of contigs

Raw nanopore signal files were first consolidated using POD5 tools, where individual POD5 files generated during sequencing were merged into a single POD5 file. The merged POD5 file was subsequently basecalled using Dorado v0.8.2 (Oxford Nanopore Technologies) with the high-accuracy sup model (dna_r10.4.1_e8.2_400bps_sup@v5.0.0). Basecalling was performed with GPU acceleration (cuda:all) to improve computational efficiency. A minimum quality score threshold of Q10 was applied to remove low-quality reads while retaining sufficient sequencing depth, which is particularly important for complex environmental samples where microbial diversity is high and rare taxa may otherwise be lost [24, 25]. Adapter trimming was disabled during basecalling (--no-trim) to avoid premature removal of sequence regions, allowing greater flexibility for downstream quality control [25]. The quality-filtered reads were saved in FASTQ format and subsequently demultiplexed using Dorado based on the SQK-RPB114-24 barcoding kit, generating barcode-resolved FASTQ files.

The reads were further processed using NanoFilt v.2.8.0 [26] to perform quality-based filtering (Q ≥ 10) and end trimming. Following quality filtering, read statistics were re-evaluated using NanoPlot v.1.43.0 [26] to assess improvements in read quality and length distributions.

Quality-filtered reads were assembled using Flye v2.9.5 [27, 28] in metagenomic mode (--meta), which is specifically designed to account for uneven coverage and high complexity typical of environmental microbial communities. The --nano-corr parameter was used to optimise assembly for corrected nanopore reads, improving contiguity and assembly accuracy. Assembly was performed with multithreading to enhance computational efficiency.

The resulting assemblies were further polished using Medaka v.2.1.1 (Oxford Nanopore Technologies), which employs neural network models to correct base-level errors in nanopore assemblies. Polishing was conducted by aligning the original reads back to the assembled contigs and generating a consensus sequence, thereby improving overall assembly accuracy. This step was repeated three times and is particularly important for nanopore-based metagenomic datasets, which are prone to higher raw read error rates compared to short-read technologies [25].

### Viral contig identification

The assembled and polished contigs were screened independently for viral signatures using three different tools: VirSorter2 [29], DeepVirFinder [30] and geNomad [31]. We used three different approaches as each tool rely on distinct sequence features and classification strategies and using multiple tools increases the robustness in identifying viral contigs. VirSorter2 uses multiple classifiers to identify diverse DNA and RNA viruses, whereas DeepVirFinder uses a convolutional neural network-based approach for alignment-free viral sequence prediction. geNomad combines gene-content information with machine-learning models for identification and annotation of viral and other mobile genetic elements [29, 30, 31].

We used VirSorter2 v2.2.4 with default settings and set a minimum contig length of 3000. The contigs with a maximum viral score (max_score) ≥0.90 were classified as high-confidence, whereas those with scores of 0.50–<0.90 were classified as moderate-confidence. The other two tools were run with default parameters and output data was categorised as per our selected criterion. For DeepVirFinder v1.0, sequences with a prediction score ≥0.90 and P < 0.05 were classified as high-confidence, while sequences with scores of 0.70–<0.90 and P < 0.05 were classified as moderate-confidence. For geNomad v1.11.1, sequences with a viral score ≥0.80, enrichment ≥1.50 and either a minimum length of 2,500 bp or at least one viral hallmark gene were classified as high-confidence. Sequences passing the geNomad default viral-score filter (score ≥0.70) but not meeting the high-confidence criteria were classified as moderate-confidence. A contig that did not meet the moderate-confidence threshold or was not reported as viral by a particular tool was assigned “None” for that tool.

To reduce tool-specific false positives and increase confidence in viral identification, predictions from the three tools were integrated into a cross-tool consensus classification. Contigs classified as high-confidence by at least two of the three tools were assigned an overall high-confidence consensus. Contigs classified as moderate-confidence by at least two tools, or as high-confidence by exactly one tool, were assigned moderate-confidence. Contigs classified as moderate-confidence by only one tool, with no additional viral prediction from the other tools, were assigned low-confidence. Contigs not flagged as viral by any tool, or receiving no confident viral prediction across the three approaches, were classified as non-viral and excluded from further analysis.

Only contigs assigned high or moderate-confidence consensus classifications were subsequently subjected to CheckV analysis to assess genome completeness and potential host contamination. CheckV v1.0.3 estimates viral genome completeness by comparison with reference viral genomes and identifies potential host-derived regions in proviral sequences. It categorizes viral sequences as complete, high quality (>90% complete), medium quality (50–90%), low quality (<50%) or undetermined when completeness cannot be estimated [32]. Medium-quality contigs were retained only when they had been assigned a high-confidence consensus classification in the preceding cross-tool analysis. Medium-quality contigs associated with a moderate-confidence consensus, as well as contigs classified by CheckV as low quality or undetermined, were excluded. The resulting dataset comprised the high-confidence, high-quality viral contigs used for downstream analyses (Figure 2).

**Figure 2:**
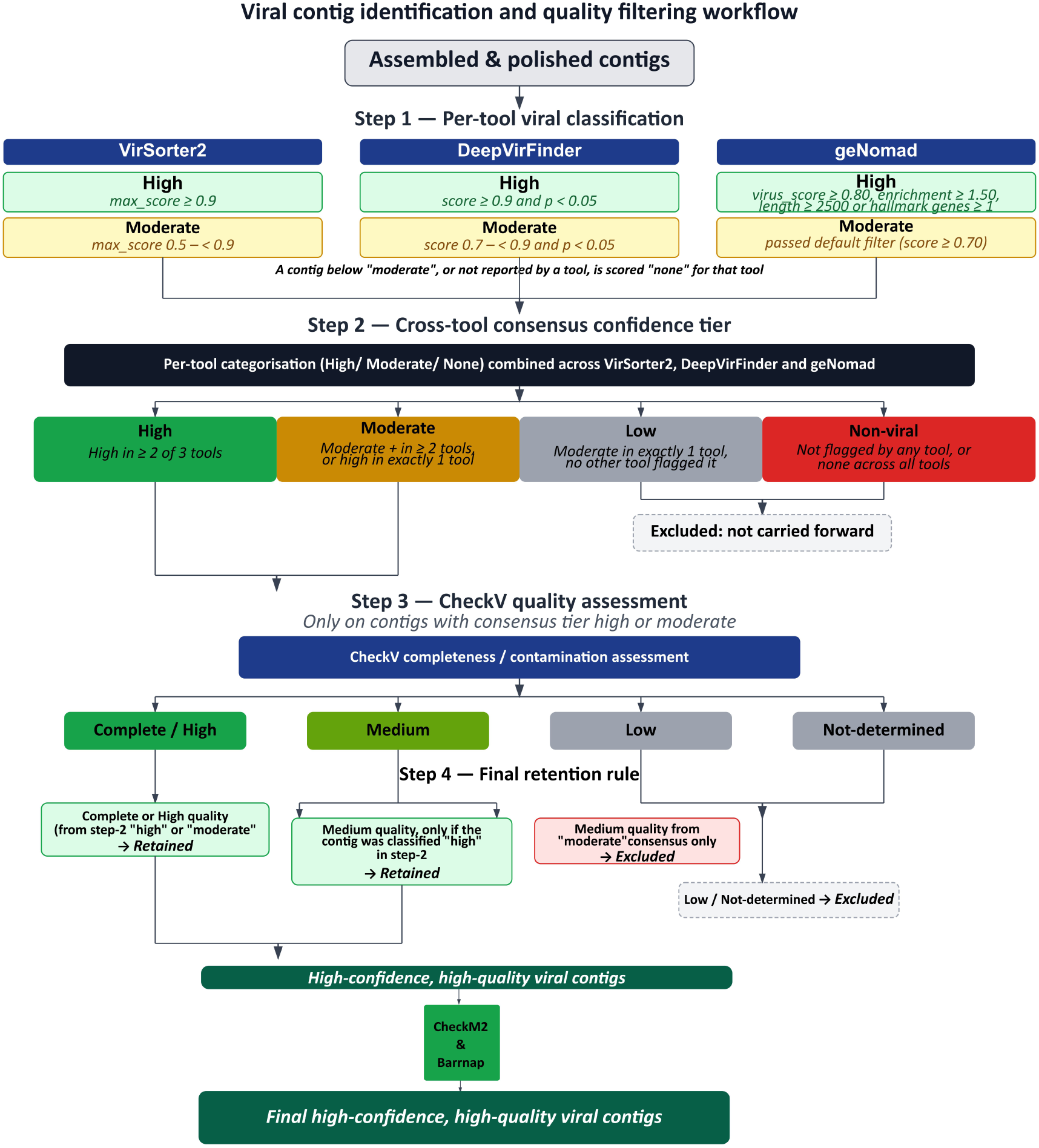
Flow-chart depicting identification of high-quality and high-confidence viral contigs among all the assembled and polished contigs. The assembled and polished contigs were classified into high and moderate confidence based on three tools: VirSorter2, DeepVirFinder and geNomad. Based on consensus of this classification, overall confidence was decided. Following this, quality of the high and medium contigs was assessed using CheckV and only the complete, high and medium quality (only for highly confident contigs) was retained.

As an additional quality check and to remove cellular contamination from the high-confidence, high-quality viral contigs, we screened for ribosomal RNA genes with Barrnap v0.9 (https://github.com/tseemann/barrnap) (bacterial HMMs, --kingdom bac, default thresholds) and for bacterial single-copy marker genes with CheckM2 v1.1.0 [33]. Contigs were removed if they contained RNA genes or had a CheckM2 completeness above 10%. In total, eight contigs were removed of 0.6-1.39 Mb, five of which carried 16S/23S/5S genes. The final high-confidence, high-quality viral contigs contained 1,604 contigs.

### Taxonomic annotation of viral contigs

Taxonomic annotation was performed using Cenote-Taker3 [34] for the 1,604 contigs. Cenote-Taker3 identifies and annotates viral genomes based on hallmark genes and curated viral databases. This approach is particularly suited for environmental viromics, as it enables detection of both known and novel viruses while minimizing false positives. Taxonomic composition of identified viral sequences was visualised using Krona [35] for Cenote-Taker3.

### Structure-informed functional annotation of complete viral contigs

Complete viral contigs with no contamination were retained after viral identification and quality assessment were subjected to genome annotation using Pharokka v1.10.0 [36]. Pharokka was used to predict coding sequences (CDSs) and other genomic features and to generate GenBank (GBK) files for downstream analysis. The resulting Pharokka-generated GenBank files were subsequently used as input for Phold v1.2.0 for structure-informed protein functional annotation. Phold integrates the ProstT5 protein language model with Foldseek structural similarity searches to identify structural homologues of predicted viral proteins [37]. Phold was executed using the default parameters, including its default annotation-confidence criteria. Phold annotations were classified according to the confidence categories reported by the software (high, medium, or low), with confidence determined from structural similarity, alignment coverage, sequence identity, ProstT5 confidence and statistical significance of the Foldseek match.

For visualization, the annotated GenBank files generated by Phold were processed using the phold plot function (Phold v1.2.0), which generates circular genome maps displaying the genomic organization and functional annotations of predicted CDS. The resulting plots were used to visualize the distribution and functional classification of annotated viral proteins along each complete viral genome.

### Viral contig clustering analysis

Viral contigs from all samples were analysed together with the vConTACT3 v3.1.4 reference viral genome database to identify gene-sharing relationships between the study-derived viral contigs and viral genomes in the database. vConTACT3 predicts protein-coding genes, identifies protein-level similarities among genomes, and integrates these relationships into genome similarity networks for viral clustering and taxonomic assignment [38].

### Host prediction analysis

Virus–host associations were inferred using two computational approaches. In the first approach, host prediction for viral genomes was performed using iPHoP v1.3.3 with default parameters and the database version iPHoP_db_Jun25_rw. The iPHoP framework integrates multiple host-association signals, including CRISPR spacer matches, sequence homology, oligonucleotide frequency similarity, and prophage-based associations, to predict putative prokaryotic hosts for viral contigs [39].

## Results & Discussion

We assembled a total of 1,604 viral contigs, of which approximately 85% were assigned at class level. Among classified viruses, *Caudoviricetes* dominated across all sampled depths. The gene sharing networks demonstrated that the viral genomes present in the database were very different from that of the ones constructed as a part of this study. This shows the limited representation of the viral genomes from the Andaman Sea and the immense diversity of viruses in the depths of oceans.

We also visualized the viral particle upon Amicon® based centrifugation and the micrographs for this can be found in Supplementary 2.

### Summary of quality of the retained assembled viral contigs

Our sequencing dataset comprised samples collected from both seamounts and transects at depths ranging from 200 m to 900 m. In total, the dataset included four transect samples and three seamount samples. The transect samples were designated as T10_ST2_200m, T11_ST3_500m, T12_ST3_900m and T13_ST1_200m, while the seamount samples were designated as SM7_ST3_500m, SM9_ST1_200m and SM9_ST3_500m. The samples were named as that “T” and “SM” denote transect and seamount samples, respectively, followed by the station number (“ST”) and the depth of sample collection.

Across all samples, sequencing depth was consistently high, with mean read counts ranging from approximately 0.6 to 3.4 million. No clear pattern was observed between the number of reads and either sample type or sampling depth. The highest and lowest mean read counts were both observed in transect samples collected at 200 m depth, but from different stations (T10_ST2 and T13_ST1, respectively).

Upon quality filtering and confidence-based scoring of the polished contigs, there were variable numbers of contigs which were retained. The checkv quality, contig length, completeness and contamination are tabulated in Supplementary file 3. The number of retained viral contigs in the samples T10_ST2_200m, T11_ST3_500m, T12_ST3_900m, T13_ST1_200m, SM7_ST3_500m, SM9_ST1_200m and SM9_ST3_500m were nine (09), six hundred seventy-two (672), forty-eight (48), eighteen (18), five hundred and twenty (520), one hundred and forty-seven (147) and two hundred (200) respectively. Most of the retained viral contigs were from high confidence consensus whereas a smaller proportion was from moderate confidence (Figure 3A).

**Figure 3:**
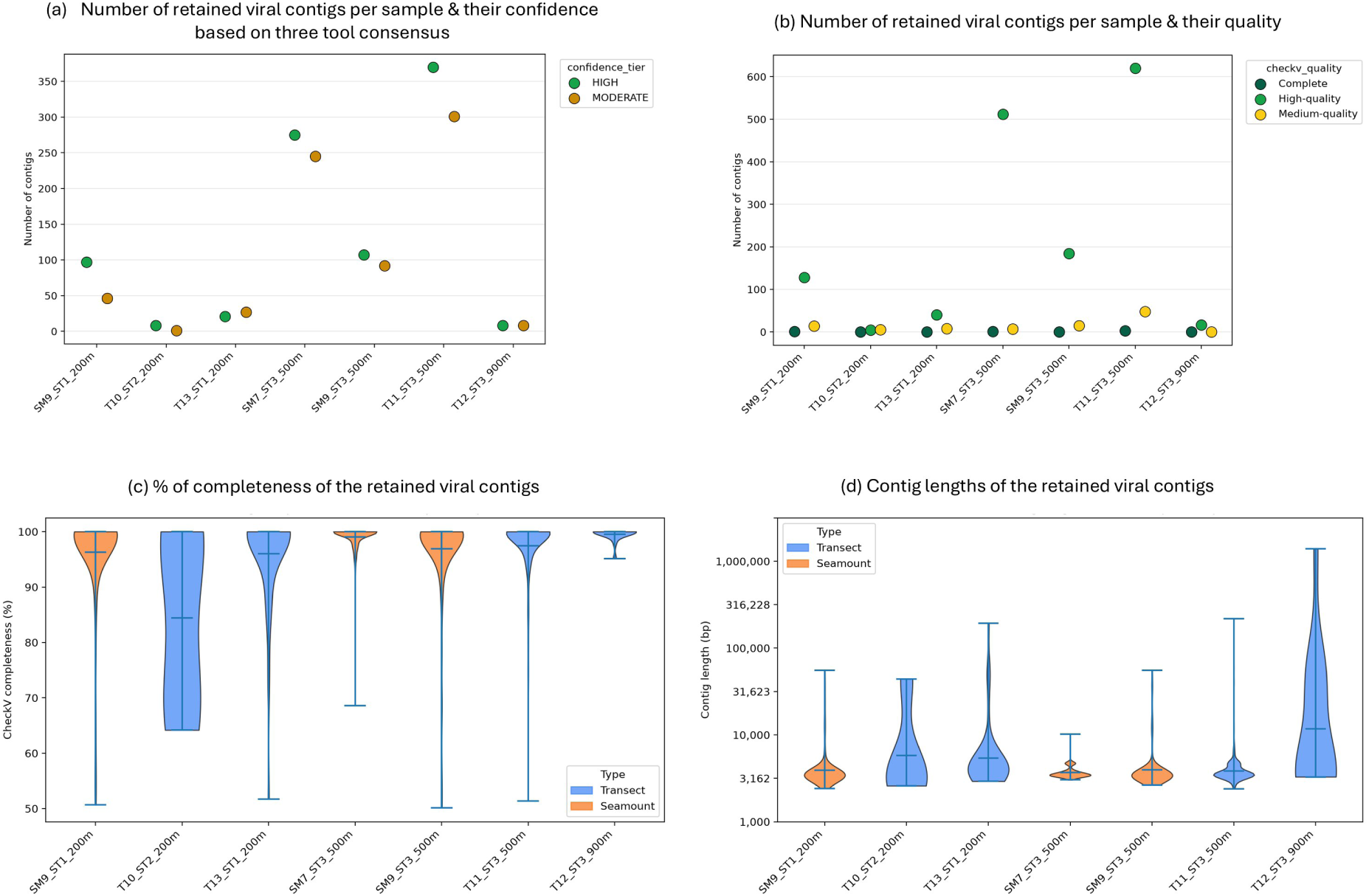
Quality statistics of retained viral contigs. (A) Number of retained viral contigs per sample and their confidence based on consensus of three tools; (B) Number of retained viral contigs per sample and their quality; (C) Percentage of completeness of retained viral contigs & (D) Contig lengths of retained viral contigs.

The final high confidence and high-quality viral contigs were predominantly of complete or high-quality (Figure 3 B). CheckV defines high-quality viral genomes as those with estimated completeness >90%, whereas medium-quality genomes have estimated completeness between 50 and 90% [32].

The distribution of estimated genome completeness provides further support for the quality of the reconstructed viral dataset (Figure 3C). Most contigs showed high estimated completeness, with the distributions concentrated near 95–100%, although one sample T10_ST2_200m contained viral contigs with a broader completeness range. The prevalence of near-complete sequences suggests that the assembly and subsequent viral-selection workflow recovered substantial portions of viral genomes rather than predominantly short genomic fragments. Long-read based sequencing approaches are advantageous with regard to obtaining near complete genomes. CheckV estimates completeness by comparing viral sequences with reference genomes and using viral protein families to derive completeness estimates for more divergent sequences.

Viral contig lengths showed a broad distribution with most sequences concentrated in the lower size range (Figure 3D). Both transect and seamount samples exhibited substantial variation in genome size, including contigs extending into the hundreds of kilobases. The occurrence of these large viral genomes is consistent with the recognized diversity of genome sizes among tailed bacteriophages (*Caudoviricetes*), which includes a distinct group of large or “jumbo” phages. Jumbo phages are commonly defined as phages with genomes >200 kb, while genomes exceeding ∼500 kb have also been described as megaphages. Previous genomic surveys have identified numerous *Caudoviricetes* genomes within these size ranges, demonstrating that exceptionally large genomes are an established feature of this viral group [40, 41]. More recent large-scale analyses have further shown that jumbo phages are widely distributed across microbial habitats, with the majority of identified jumbo phage genomes belonging to *Caudoviricetes* and genome sizes extending beyond 800 kb [42]. Thus, the long contigs recovered in the present study may represent members of the large-genome phage fraction rather than being unexpected outliers in viral genome size. Nevertheless, genome size alone was not used as a criterion for viral quality or retention and the presence of large contigs should be interpreted in conjunction with their CheckV completeness and contamination estimates. The large genomes also showed no contamination as per CheckV results (Supplymentary file 3).

### Genome size distribution of the retained assembled viral contigs

The final high-confidence, high-quality viral contigs showed a right-skewed distribution of genome lengths across all samples (Figure 4A). The density distributions showed that the majority of contigs were concentrated within the ∼3–6 kb range. However, all samples also displayed extended right-hand tails, indicating the presence of a smaller fraction of substantially larger viral contigs (Figure 4B). The extent of this long-tail distribution varied among samples, with some samples showing a considerably broader genome-size distribution than others. Thus, although the retained viral dataset was dominated by relatively short contigs, a comparatively small subset represented exceptionally large viral genomes.

**Figure 4:**
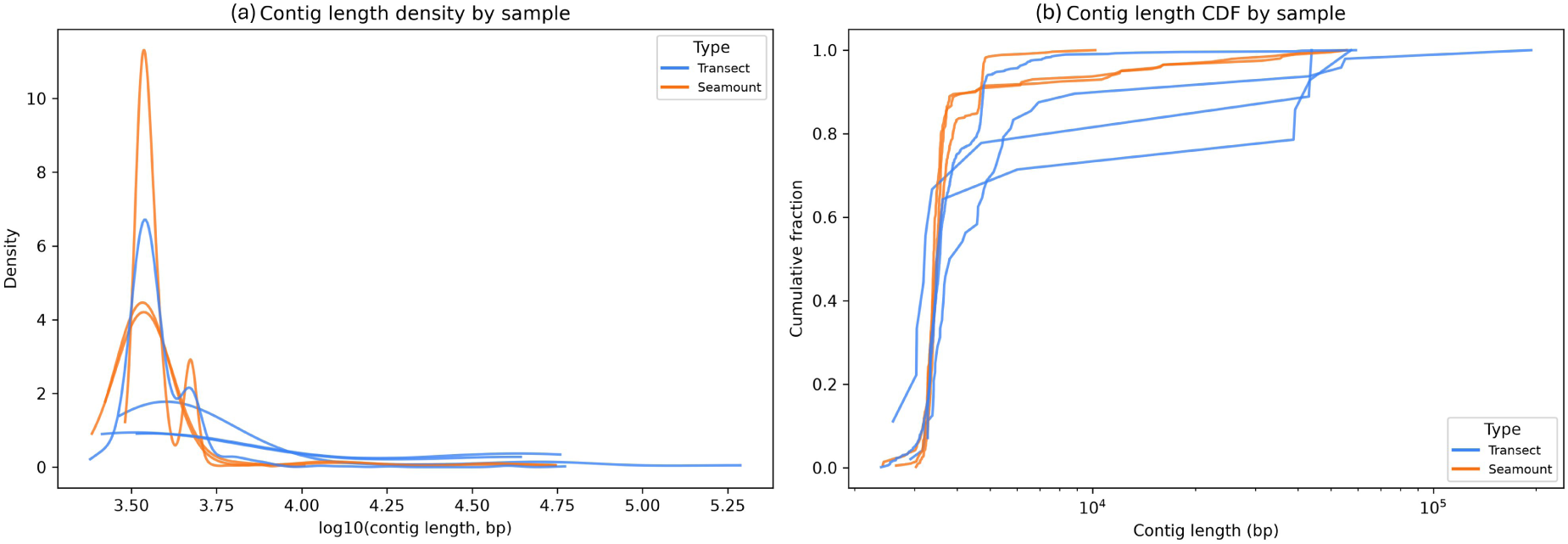
Length of retained viral contigs. (A) Plot for density distributions of viral contig lengths plotted as log10 (contig length), showing more contigs in the lower kilobase with long-tailed distributions extending toward larger contigs. (B) Cumulative distribution of contig length plotted on a logarithmic (x-axis). Most viral contigs occurred below approximately 10 kb, although all samples contained a smaller fraction of substantially longer contigs, with the longest contigs extending into the hundreds of kilobases.

The occurrence of these large genomes is biologically plausible, particularly given the increasing recognition of jumbo and huge bacteriophages in metagenomic datasets. Tailed bacteriophages with genomes >200 kb is generally referred to as jumbo phages, and environmental metagenomic studies have revealed that such large phages are much more widespread than suggested by earlier culture-dependent surveys [40, 41].

Importantly, the presence of exceptionally long contigs should not be interpreted solely on the basis of length. Large assembled sequences can potentially arise from assembly-related artefacts, including repeat-induced mis-assembly or concatenation of genomic regions. Such issues have previously been documented for metagenomically reconstructed jumbo phages, emphasizing the importance of independent completeness, contamination and assembly-quality assessments when interpreting exceptionally large viral contigs [43]. In the present study, the large contigs considered here had already passed the high-confidence viral-identification and high-quality CheckV filtering criteria, providing additional support for their consideration as candidate large viral genomes.

### Taxonomic assignments of viral contigs

Cenote-Taker3 provided taxonomic assignments for a substantial proportion of the retained viral contigs, although the proportion of sequences assigned decreased with increasing taxonomic resolution (Figure 5). At the class level, 88–100% of viral contigs were assigned across the seven samples, whereas assignment rates were progressively lower at the order and family levels (Figure 5B). This pattern indicates that the recovered viruses could generally be placed within broad viral taxonomic groups, while a substantial fraction could not be resolved to lower taxonomic ranks. Such a decline in taxonomic resolution is expected for metagenome derived viral sequences, particularly those originating from poorly represented or highly divergent viral lineages. This is because taxonomic assignment depends on the availability of homologous viral sequences and conserved viral marker genes in reference databases. Cenote-Taker3 is specifically designed to provide hierarchical viral taxonomy using hallmark genes and reference taxonomic databases, including for divergent viral sequences [34].

**Figure 5:**
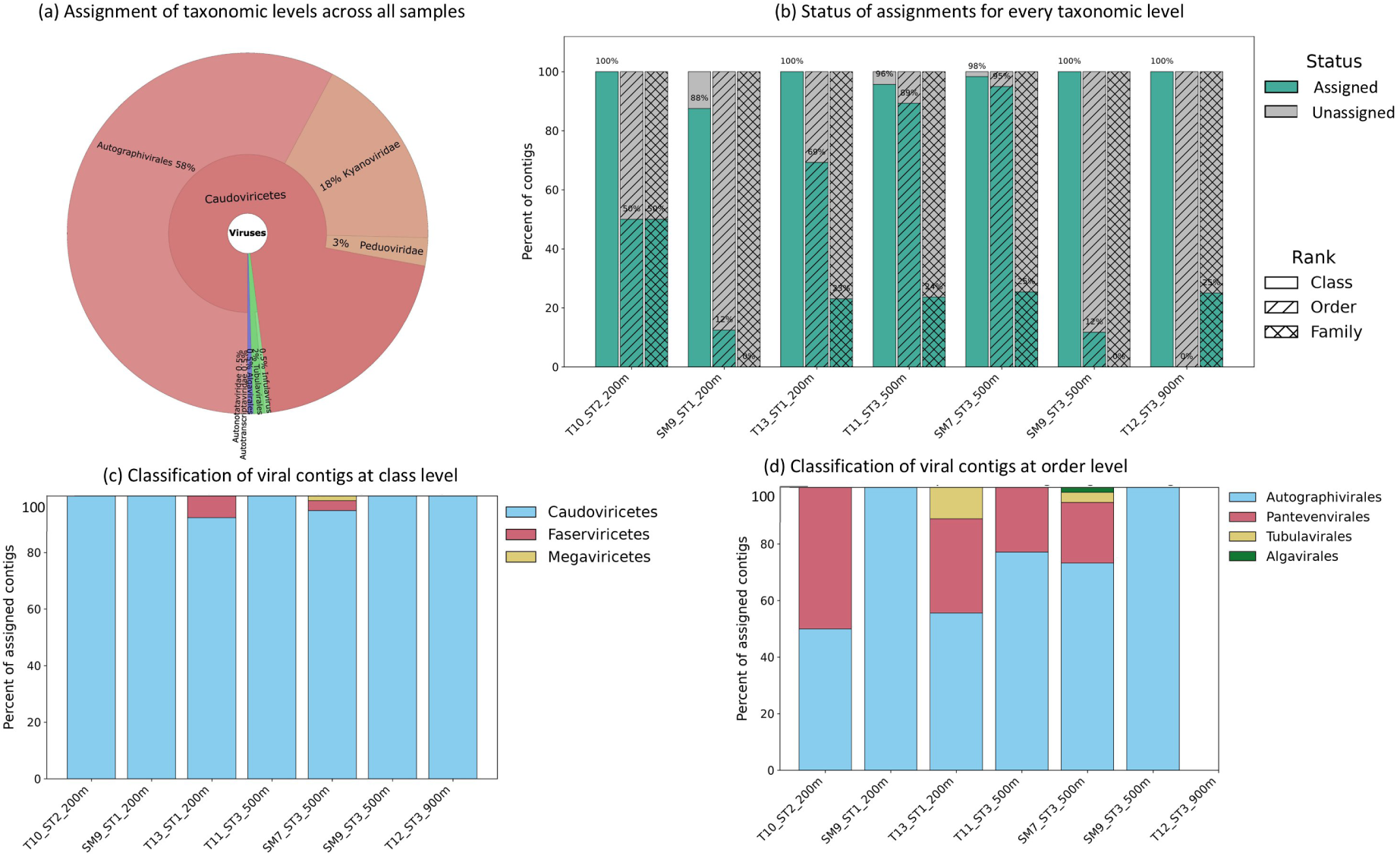
Taxonomic composition of retained viral contigs inferred using Cenote-Taker3 based on detection of viral hallmark genes. (A) Overall distribution of taxonomic assignments across the retained viral contigs, showing the hierarchical classification from class to lower taxonomic ranks. (B) Percentage of viral contigs assigned and unassigned at the class, order and family levels for each sample. Assignment rates generally decreased with increasing taxonomic resolution with a larger proportion of contigs remaining unassigned at the family level. (C) Relative composition of assigned viral contigs at the class level for each sample, showing the predominance of Caudoviricetes and comparatively minor contributions from Faserviricetes and Megaviricetes. (D) Relative composition of assigned contigs at the order level.

Across the assigned contigs, *Caudoviricetes* was overwhelmingly the dominant class, accounting for approximately 92–100% of class-level assignments in most samples (Figure 5C). Only a small proportion of assigned contigs belonged to other known classes, including *Faserviricetes* and *Megaviricetes*. The dominance of *Caudoviricetes* was also reflected in the order-level composition, although the relative representation of individual orders varied among samples (Figure 5D). These differences suggest that the viral communities recovered from the different sampling locations were not taxonomically uniform.

The high representation of *Caudoviricetes* is consistent with previous metagenomic surveys showing that tailed bacteriophages are frequently among the dominant dsDNA viruses recovered from aquatic environments, including marine, estuarine and freshwater ecosystems [44]. However, this pattern should be interpreted in the context of reference-database composition and taxonomic assignment bias. Viral reference databases remain disproportionately populated by characterized bacteriophages relative to the much larger diversity of uncultivated environmental viruses, which can increase the probability that environmental viral sequences are assigned to *Caudoviricetes*. Indeed, previous studies have explicitly noted that the apparent dominance of *Caudoviricetes* in metagenomic datasets may partly reflect their overrepresentation in available reference databases [45]. Conversely, a substantial fraction of environmental viral sequences often remains unclassified, particularly at lower taxonomic ranks, because of limited representation of closely related reference genomes and the high sequence divergence of uncultivated viruses [46]. Thus, the progressively lower assignment rates observed at order and family levels in the present study may reflect both limited taxonomic resolution of current viral databases and the presence of highly divergent or previously underrepresented viral lineages.

### Functional annotation of high-confidence & high-quality viral contigs

All of the high-confidence and high-quality viral contigs were further examined using Phold, a structure-informed phage annotation framework against a curated database of >1.36 million phage protein structures [37]. Three viral contigs were categorized by CheckV as “complete”. All three genomes were approximately 4.9 kb in length and were classified as complete by CheckV and with no detectable contamination (Figure 6).

**Figure 6:**
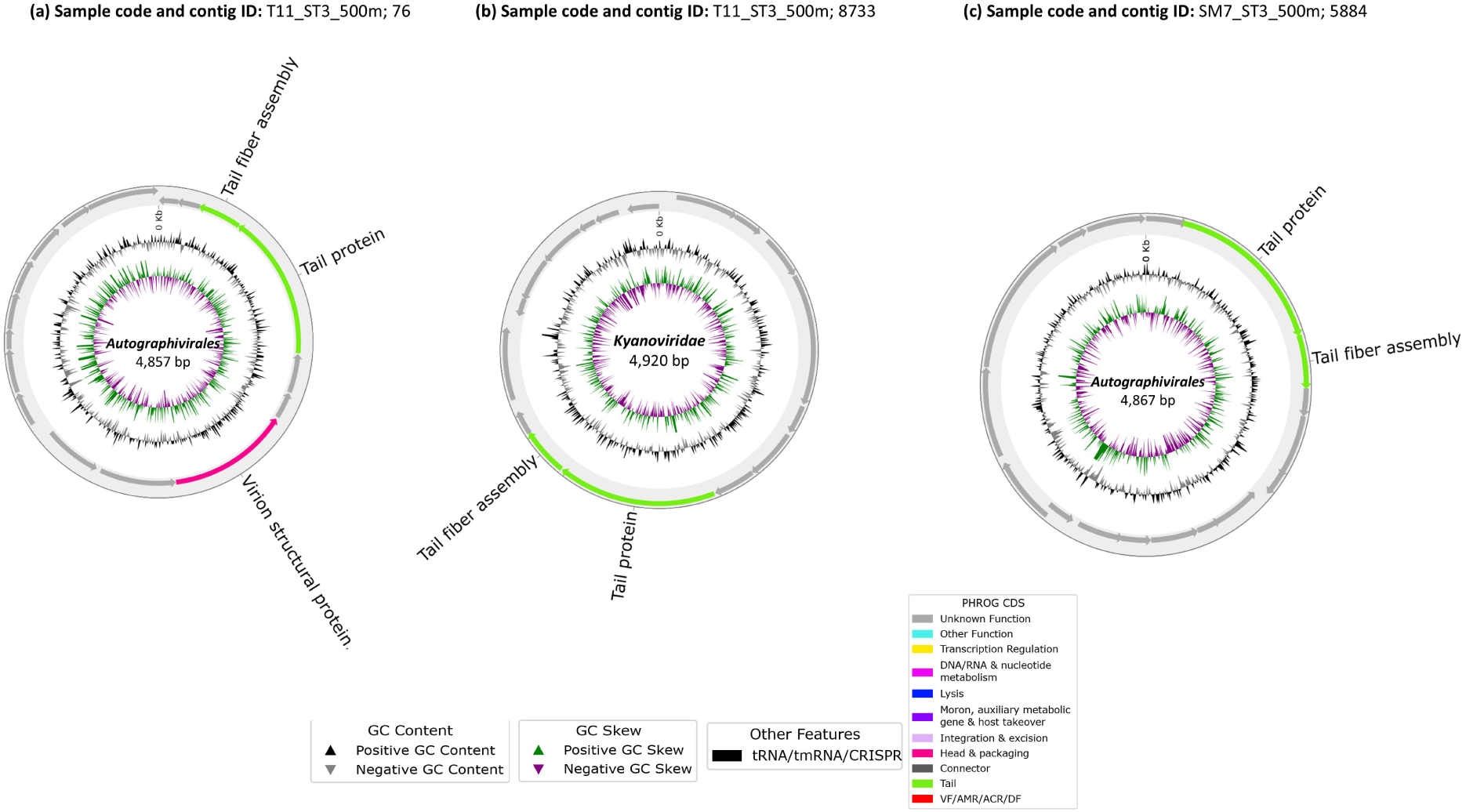
Genome organization and structure-informed functional annotation of three complete viral genomes of approximately 4.9 kbp. Circular genome maps show the genomic organization and predicted protein functions of three representative complete viral genomes. All three genomes were classified as Caudoviricetes and showed 100% completeness and 0% estimated contamination according to CheckV. The genomes shown are T11_ST3_500m;76 (4,857 bp), classified within the order Autographivirales; T11_ST3_500m;8733 (4,920 bp), classified within Kyanoviridae; and SM7_ST3_500m;5884 (4,867 bp), classified within the order Autographivirales.

Each genome contained 17 predicted coding sequences (CDSs). Phold generated functional predictions for only a subset of these CDSs, with annotation percentage increasing from 2/17 CDSs (11.8%) in SM7_ST3_500m (contig_5884), to 3/17 (17.6%) in T11_ST3_500m (contig_8733) and 5/17 (29.4%) in T11_ST3_500m (contig_76). Thus, despite the complete status of these genomes, 70.6–88.2% of predicted proteins remained without a Phold functional assignment, illustrating the substantial degree of functional dark matter even within complete environmental viral genomes. All the three viral contigs were annotated for tail associated functions. A tail-fibre assembly protein (PHROG 4912) was identified with high confidence in all three genomes, with sequence-structure matches supported by high Foldseek similarity. Several other proteins produced structural matches to PHROGs but remained classified as unknown function.

The predominance of tail-associated proteins is consistent with the genomic architecture expected for small tailed bacteriophages. In particular, the identification of a tail-fibre assembly protein across all three genomes provides independent structural evidence supporting their classification as *Caudoviricetes* and indicates conservation of host-interaction machinery despite their environmental source.

Apart from the three complete viral contigs, we selected six high-confidence viral contigs with genome size approximately 40-50 kbp for representation of a detailed genomic annotation for functions using Phold (Figure 7). The contigs ranged from 39,244 to 59,251 bp in length and were classified as members of the class *Caudoviricetes* by both Cenote-Taker 3 and geNomad. CheckV estimated genome completeness of 100% (except for SM9_ST3_500m with 99.89% completeness) and had no contamination.

**Figure 7:**
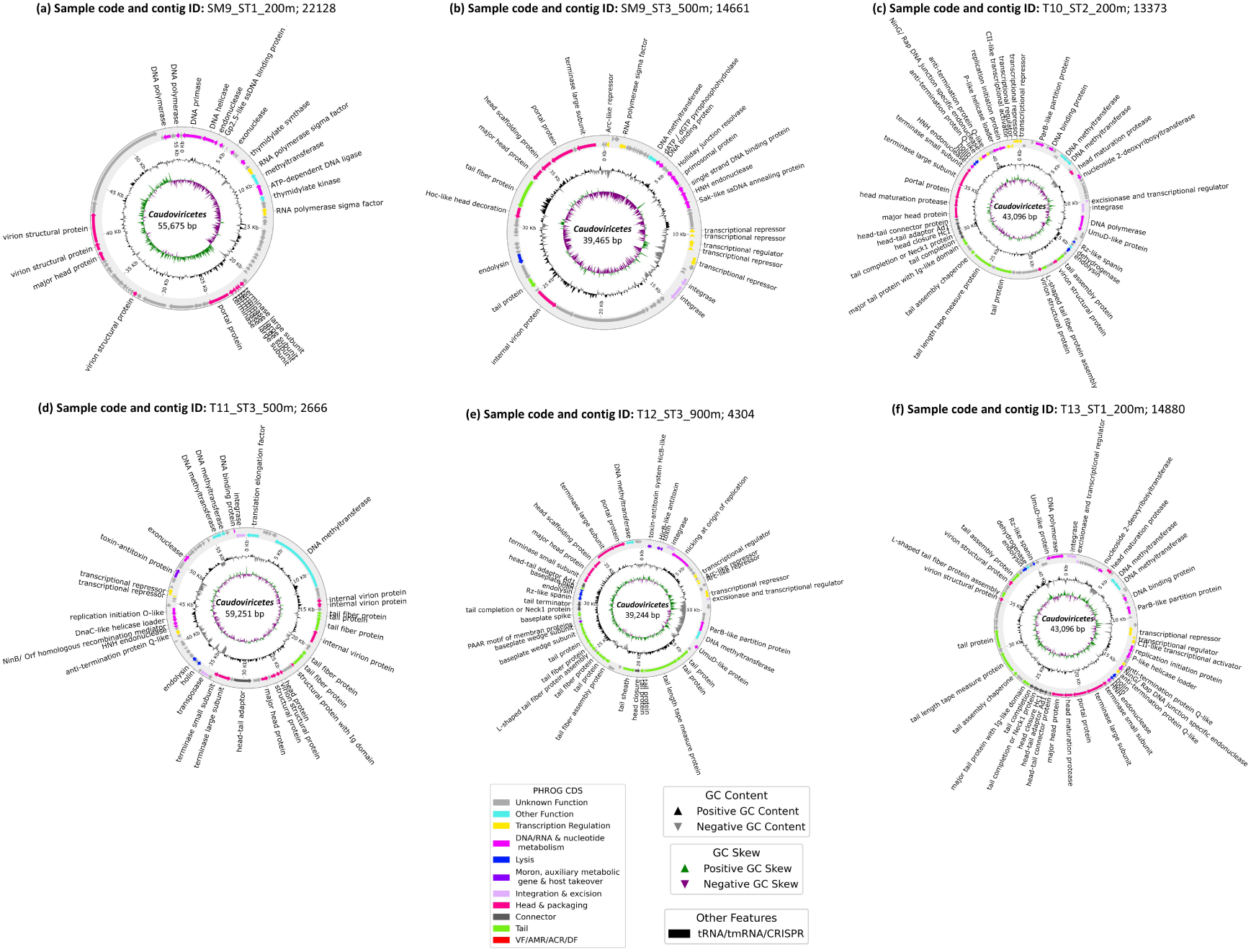
Genome organization and structure-informed functional annotation of six high-quality viral genomes of 40-50 kbp length. Panels (a–f) are representative viral contigs reconstructed from SM9_ST1_200m (55,675 bp), SM9_ST3_500m (39,465 bp), T10_ST2_200m (43,096 bp), T11_ST3_500m (59,251 bp), T12_ST3_900m (39,244 bp) and T13_ST1_200m (43,096 bp) respectively. Predicted coding sequences (CDSs) are annotated using Phold. The inner tracks represent GC content and GC skew, with positive and negative values indicated by triangles of different orientations and colours, as shown in the legend. All of the viral contigs were assigned to Caudoviricetes by Cenote-Taker3 and geNomad.

Phold annotation revealed a diverse repertoire of predicted proteins associated with viral genome replication, transcriptional regulation, virion assembly, DNA packaging, recombination, and host–virus interactions (Figure 7). Structural and assembly-related proteins included major head or capsid proteins, portal proteins, head-maturation proteases, tail proteins, tail-fibre proteins, tail assembly proteins and terminase large subunits. These annotations were particularly prominent in the six genomes, supporting their identification as tailed bacteriophages. The occurrence of head, portal, tail, and packaging-associated proteins is consistent with the coordinated assembly and genome-packaging machinery characteristic of many members of *Caudoviricetes* [47]. In tailed bacteriophages, the portal protein forms a vertex of the capsid and provides a channel for DNA translocation, while the terminase complex recognizes and processes viral DNA and powers its packaging into the capsid. Tail-associated proteins subsequently contribute to virion assembly and interactions with susceptible hosts [48].

Proteins associated with genome replication and maintenance were also predicted across the contigs. These included DNA polymerases, DNA helicases, primase-related proteins, DNA methyltransferases, ATP-dependent DNA ligases, endonucleases and recombination-associated proteins. Additional annotations included single-stranded DNA-binding proteins and proteins associated with DNA repair or processing. Collectively, these predicted functions indicate that the genomes encode components potentially involved in viral DNA synthesis, processing and maintenance during infection. However, the precise replication mechanisms cannot be established from these annotations alone. The combination of structural and replication-associated genes is consistent with the multifunctional organization of tailed bacteriophage genomes, in which distinct but sometimes overlapping functional modules support genome propagation and virion production [49].

Transcriptional and regulatory proteins formed another prominent functional category. These included RNA polymerase sigma factors, transcriptional repressors, transcriptional regulators, transcriptional activators and anti-termination-associated proteins. Such proteins are involved in diverse mechanisms of bacteriophage transcriptional regulation, including modulation of host RNA polymerase activity and control of transcription initiation and termination [50, 51]. The occurrence of integrases, excisionases and recombination-related proteins in some genomes further indicates the potential for genome integration or recombination, although these functions do not independently establish a lysogenic lifestyle.

In temperate bacteriophages, integrases and excisionases or recombination directionality factors can mediate prophage integration and excision, nevertheless, the presence of these genes alone does not establish a lysogenic lifestyle [52, 53].

Other predicted proteins included nucleotide-metabolism enzymes, nucleoside-diphosphate kinases, nucleotide kinases and proteins associated with translation or protein modification. Several genomes also contained annotations related to partitioning, chromosome segregation or DNA-binding functions. A substantial proportion of predicted coding sequences were assigned to unknown or broadly defined functions, highlighting the limitations of current databases in characterizing uncultivated marine viruses. Phold is particularly useful for investigating such genomes because it uses protein structure-informed annotation to identify potential functions that may not be readily detected by conventional sequence-similarity searches. This approach improves annotation sensitivity for divergent phage proteins, although predicted structural similarity does not by itself demonstrate biochemical activity [37]. This limitation is particularly relevant to metagenomic studies of marine viruses, where the majority of viral diversity is not represented by cultured reference genomes. Large-scale marine virome surveys have repeatedly revealed extensive viral sequence diversity and substantial numbers of previously uncharacterized lineages [54]. The reconstruction of the highly complete *Caudoviricetes* genomes from deep-ocean samples extends the genomic information available for viruses present in the Andaman Sea.

### Gene sharing relationships and genomic novelty of reconstructed high-confidence and high-quality viral contigs

We found that many of the viral contigs remained unassigned at lower taxonomic rank in Cenote-Taker3 and functionally unannotated in Phold. Hence, comparing these assembled contigs with the publicly available genomes would provide an idea about their genomic relatedness. To understand the genomic relatedness of the 1,612 retained viral contigs from this study, we used vConTACT3 and analyzed together with reference viral genomes. The network revealed extensive genomic diversity among the viral contigs from our study (Figure 8). Among the 1,612 viral contigs, only 32 received a named class-level assignment, comprising 30 contigs assigned to *Caudoviricetes* and two to *Faserviricetes*. At lower taxonomic ranks, assignment to known taxa was considerably more limited, with only two contigs assigned to order *Tubulavirales*, four to families *Peduoviridae* and *Inoviridae*, and one to a genus *Infulavirus*. This limited assignment to established taxa indicates that most of the recovered sequences occupy genomic space that is poorly represented by currently available reference viral genomes. Gene sharing networks showed that the 1,604 viral contigs were distributed among 22 novel order-level groups, 40 novel family-level groups, 63 novel subfamily-level groups and 699 novel genus-level groups. Thus, the high number of novel network-defined groups indicates substantial genomic heterogeneity among the recovered viruses and suggests that the dataset captures viral diversity that extends beyond currently characterized lineages.

**Figure 8:**
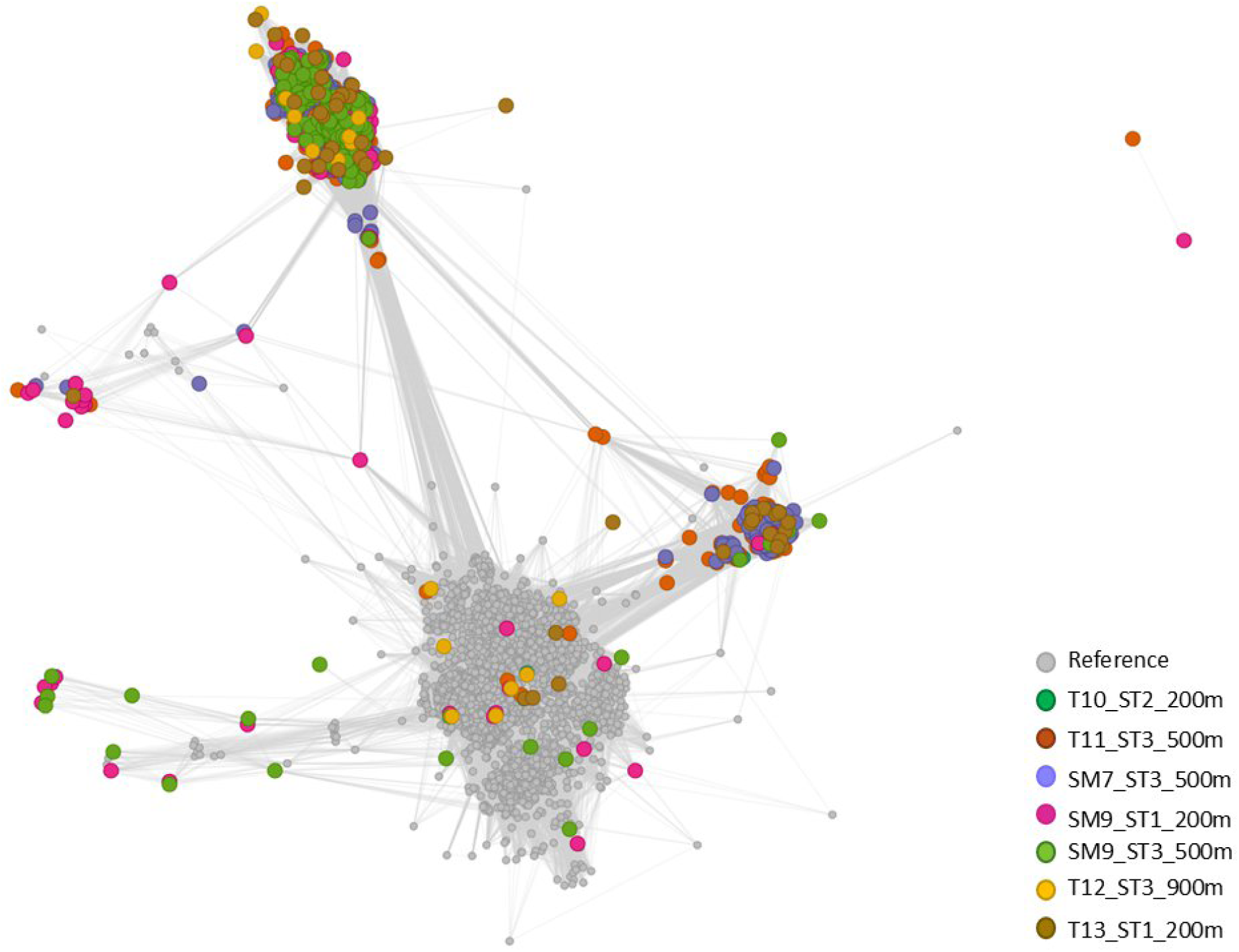
Gene-sharing network of viral contigs and reference viral genomes. The network was generated using vConTACT3 based on shared protein content among viral genomes. Each node represents a viral genome or contig, with grey nodes representing reference viral genomes and colored nodes representing viral contigs reconstructed in this study. Colours indicate the corresponding sampling location.

Some of the 1,604 viral contigs from this study were also found to occupy the clusters containing reference genomes indicates that some of the viruses share detectable protein-level genomic relationships with previously characterized viruses. In contrast, most viral contigs from our study, occupied more peripheral regions of the network or forming groups with limited representation of reference genomes showed comparatively weak gene-sharing relationships with known viruses. This pattern suggests a continuum of genomic relatedness, ranging from viruses with characterized lineages to highly divergent viral groups with limited representation in existing databases.

The extensive representation of novel groups of viral contigs is consistent with the growing recognition that environmental viral communities contain a large fraction of uncultivated and poorly characterized genomic diversity. This is particularly true for the deep ocean virome which is largely unexplored. Gene-sharing approaches are particularly useful in this context because viral genomes can be highly divergent at the nucleotide level while retaining subsets of homologous proteins that allow their relationships to be detected. vConTACT3 integrates protein-clustering information to establish genome-level relationships and hierarchical taxonomic predictions, allowing viral sequences to be placed within a broader genomic context even when conventional taxonomic assignment is not possible [38].

The limited representation of known taxa and the genomic heterogeneity reflect a combination of incomplete representation of viral diversity from oceans and the highly divergent viruses present in such ecosystems.

### Predicted virus-host relationships in the oceanic deep waters

Potential hosts of the recovered viral contigs were inferred using iPHoP (integrated Phage–Host Prediction), which combines complementary virus- and host-based approaches to predict hosts of bacteriophages and archaeal viruses. iPHoP integrates multiple prediction methods using a machine-learning framework and is designed primarily for host prediction at the genus level while maintaining a low estimated false-discovery rate [39].

Among the 1,612 retained viral contigs, a host was predicted in 10.7% of them (Figure 9). The predicted host associations were mostly bacterial (98.3%) and rest were archaeal (1.7%). At the host-phylum level, Pseudomonadota represented the predominant predicted host group, followed by smaller contributions from Bacillota, Thermoplasmatota and Chloroflexota. The high proportion of unassigned hosts should not necessarily be interpreted as evidence that these viruses lack hosts. Rather, it may reflect the limited representation of their actual or closely related host genomes in available reference databases.

**Figure 9:**
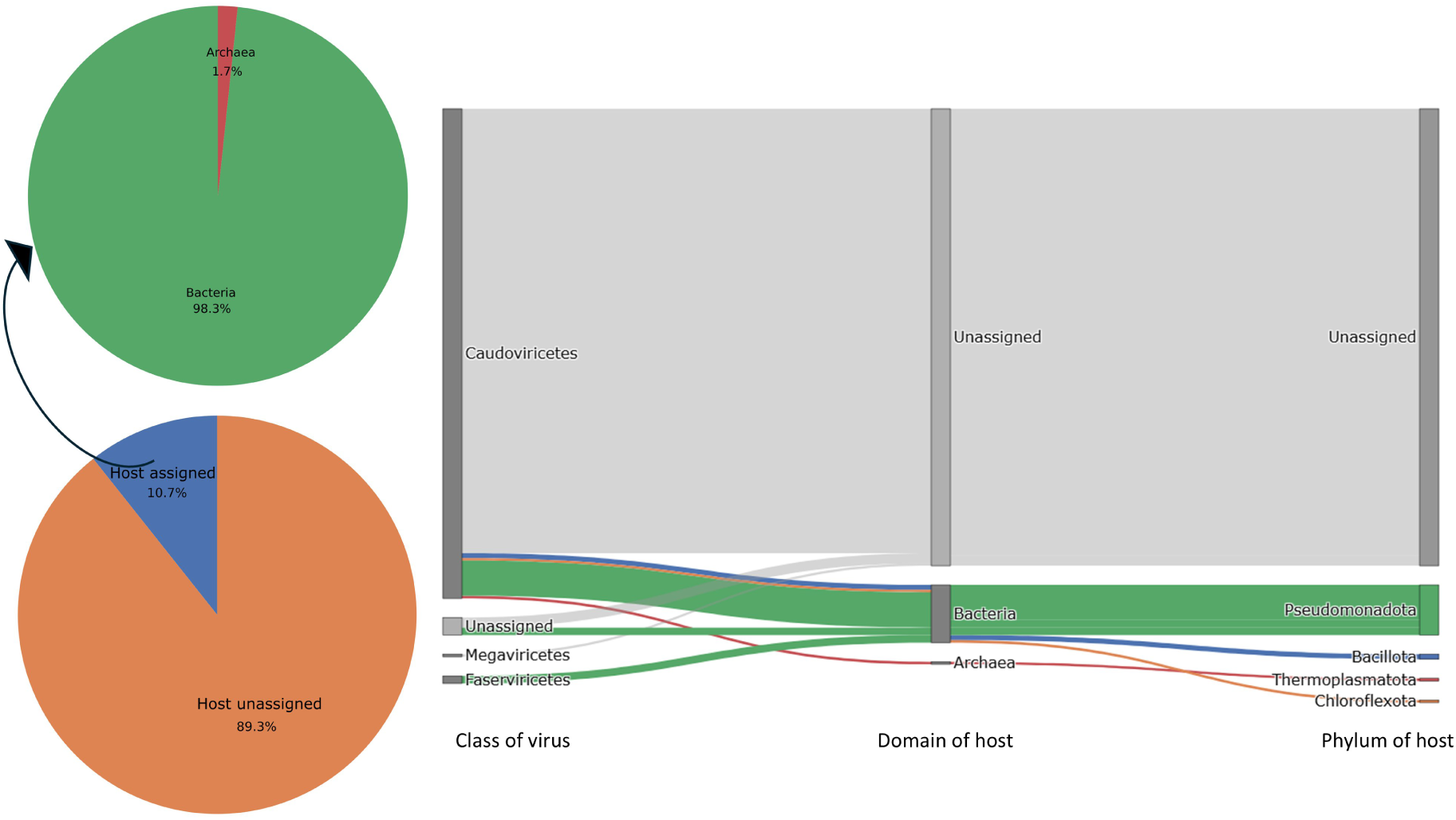
Predicted virus–host associations based on iPHoP. The lower-left pie chart shows the proportion of viral contigs for which a host association was predicted versus those remaining unassigned. Among viral contigs with predicted hosts, the upper-left pie chart shows the inferred host domain composition, with Bacteria accounting for 98.3% and Archaea for 1.7% of predicted hosts. The Sankey diagram illustrates the hierarchical relationships between viral class, predicted host domain and host phylum.

Among the assigned bacterial host phylum, the presence of Pseudomonadota was highest fraction of it. This might be because Pseudomonadota (formerly Proteobacteria) is one of the most diverse and abundant bacterial phyla in aquatic environments, with major lineages occurring across both marine and freshwater bacterioplankton communities [55,56]. However, because iPHoP predictions are computational inferences, these associations should be interpreted as putative virus–host linkages rather than experimentally validated outcomes.

## Conclusion and future directions

This study provides the first comprehensive characterization of communities of DNA viruses from the mesopelagic waters of the Andaman Sea using long-read metagenomic sequencing. We used three complementary approaches to identify high-confidence and high-quality viral contigs amongst thousands of assembled and polished contigs. This yielded 1,604 such viral contigs. Taxonomic analyses revealed that a large proportion of the recovered viral sequences had limited representation among currently characterized viral genomes. Although *Caudoviricetes* represented the dominant assigned viral class, gene-sharing analysis revealed extensive representation of novel groups across multiple taxonomic levels. This pattern highlights the substantial underrepresentation of environmental viruses in reference databases. Host prediction further indicated that most confidently predicted virus–host associations involved bacteria, with Pseudomonadota representing the predominant predicted host phylum. However, host associations could be inferred for only a relatively small proportion of the recovered viral sequences. Functional analysis of viral genomes also illustrated the extent of viral functional dark matter. This is not unexpected for environmental viral genomes, for which sequence-based homology often provides limited functional information.

While this study establishes a baseline for mesopelagic viral ecology in the Andaman Sea, several important questions remain that will require future studies. The large proportion of unclassified viral sequences highlights the need for expanded marine viral reference databases, particularly from understudied regions such as the Indian Ocean. Second, this study was restricted to DNA viruses, but the RNA viruses also remain entirely understudied in this region. Third, integration of meta-transcriptomic data with metagenomic datasets would enable identification of AMGs that are actively expressed and the environmental conditions favoring their expression.

As the oceans continue to warm and stratify under accelerating climate change, understanding the viral communities that regulate deep-sea microbial processes will be essential for predicting the future of ocean biogeochemistry.

## Acknowledgements

We would like to thank Centre for Marine Living Resources and Ecology (CMLRE), Kochi, Kerala, India and National Institute of Ocean Technology (NIOT), Chennai, Tamil Nadu, India for the logistical assistance for sampling.

## CRediT authorship contribution statement

**Manisha Ray:** Conceptualization, Formal analysis, Investigation, Methodology, Software, Validation, Visualization, Writing – original draft, Writing – review & editing. **Muthuvarmadam Subramanian Ram**: Conceptualization, Data curation, Methodology, Writing – review & editing, **Himanshu Kothari:** Data curation, Methodology. **Mamidala Srilakshmi:** Methodology, **Harinivas Harshan Krishnan:** Conceptualization, Funding acquisition, Writing – review & editing. **Govindhaswamy Umapathy:** Conceptualization, Funding acquisition, Project administration, Supervision, Writing – review & editing.

## Author Notes

Three supplementary files are available with the online version of this article.

**Supplementary file 1:** Details of all the sampled points including their date of sampling, volume of water filtered, geocoordinates and abiotic metadata.

**Supplementary file 2:** Transmission Electron Microscopic images showing viral particles isolated from deep ocean water sample upon Amicon® centrifugation.

**Supplementary file 3:** Details (contig length, completeness, contamination) of the high-quality and high-confidence viral contigs retained and analyzed in this study.

## Abbreviations

IPCC: Intergovernmental Panel on Climate Change
AMGs: auxiliary metabolic genes
GOV: Global Ocean Virome
TFF: tangential flow filtration
CTD: conductivity–temperature–depth
PAR: photosynthetically active radiation
PC: polycarbonate
TEM: Transmission Electron Microscope
ONT: Oxford Nanopore Technologies
PHROGs: Prokaryotic Virus Remote Homologous Groups
CDS: Coding Sequence

## Impact Statement

Marine viruses are major components of ocean ecosystems, yet their genomic diversity remains poorly characterized. This is particularly true for the Indian Ocean particularly at depths of these oceans. This study presents the first comprehensive dataset of the virome of the mesopelagic region of the Andaman Sea. We reconstructed 1,604 high-confidence and high-quality viral contigs using long-read metagenomics. This unveiled substantial dark-matter and reveled the genomic novelty of these viruses. This study expands the genomic representation of the viruses from the deep ocean and provides a valuable resource for the understanding of their evolution, host interactions and ecological roles.

## Data Summary

The raw sequencing data generated in this study have been deposited in the NCBI Sequence Read Archive (SRA) under BioProject accession PRJNA1466812. The dataset has seven samples with SRR IDs from SRR38678299–SRR38678305, corresponding to BioSample accessions SAMN60171981–SAMN60171987. All sequencing data have been released and are publicly accessible through the NCBI SRA database. The final high-quality and high-confidence viral contigs reconstructed as a part of this study is also submitted in the same BioProject.

All the code and commands to run the bioinformatic tools are publicly available.

